# A context-aware analytical framework for DNA methylation biomarker discovery

**DOI:** 10.64898/2026.02.20.706794

**Authors:** Agustina Renata Angeloni, Clara García Samartino, Juan Sebastián Arbona, Cintia Celina Vaquer, Paula Alida Wetten, Victoria Bocanegra, Rodrigo Damián Militello, Guillermo Sanguinetti, Agustín Correa, Melani Carlen, Walter Ramon Minatti, Gastón Andres Vaschalde, Roberto Perez Ravier, Ricardo Nicolás Manzino, Jorge Daniel Rodriguez, Paula Valdemoros, Leandro Sarrio, Alejandro Ledesma, Emanuel Martín Campoy

## Abstract

DNA methylation biomarker discovery commonly begins with tumor-associated differences at individual CpGs or genomic regions, yet the utility of a candidate depends on how its signal behaves across the biological backgrounds encountered in its intended application. Reference tissues, alternative tumor backgrounds and cohort composition can therefore shape whether a methylation change is informative. Here we present a gene-centered framework that integrates methylation contrast, intersample variability, context-specific methylation reference layers and sequence features to prioritize coordinate-defined genomic intervals. Applied to colorectal, hepatocellular and lung cancers, the framework revealed context-dependent shifts in candidate selection and regional metrics. Selected regions and a benchmark locus were then translated into native-DNA quantitative PCR assays that preserved discriminatory methylation signals in independent colorectal and liver tissue cohorts. Together, these results establish a context-aware route from genome-scale methylation data to experimentally addressable biomarker targets.

## Introduction

DNA methylation profiling captures reproducible epigenetic programs that distinguish cell types and enable inference of cellular origin from bulk methylomes ^1,2^. In cancer, these patterns retain clinically relevant information across tumor tissue and circulating DNA, supporting molecular subtyping ^3^, multicancer detection, including early-stage disease ^4^, and classification of tissue of origin across diverse tumor types ^5^. Pan-cancer analyses have further identified differentially methylated regions that are either shared across malignancies or associated with particular tumor types, histological subtypes or tissues of origin, supporting regional methylation patterns as informative units for biomarker discovery ^6^.

Despite this potential, the biomarker value of a methylation signal depends on the biological background, cellular composition and disease context against which it is evaluated. The relevance of a candidate methylation signal can also depend on the cellular composition and disease context of the biological specimen. Methylation-based deconvolution of plasma has shown that leukocytes contribute most circulating cell-free DNA, including in patients with solid tumors, whereas DNA released by tumor cells represents a smaller and variable fraction ^7^. This provides one illustration of how biological background can influence the usefulness of a candidate methylation signal. A prospective CRC screening study has likewise shown markedly different detection performance for invasive cancer and advanced precancerous lesions, illustrating that biomarker performance depends on the biological state targeted by the intended application ^8^. Biomarker performance also changes with the reference states included during development. Recent hepatocellular carcinoma studies have therefore evaluated methylation markers against chronic hepatitis, cirrhosis, other liver conditions associated with elevated cancer risk and non-target malignancies ^9,10^. More broadly, classifiers trained across platforms can distinguish more than 170 tumor entities from their methylation profiles ^11^, demonstrating that tumor methylomes retain information beyond a binary comparison with normal tissue ^3^. Collectively, these findings indicate that biomarker discovery must be aligned with the biological comparisons required by the intended application.

Current approaches capture complementary components of methylation biomarker development. Large-scale resources provide coordinate-resolved catalogues of cancer-associated differentially methylated regions and their genomic annotations ^12^. Multi-omic analyses can further connect methylation changes with gene expression and regulatory activity ^13^. Recent computational frameworks have also addressed candidate reliability by accounting for measurement noise, individual characteristics and potentially causal disease associations ^14^. Targeted sequencing studies have shown that genome-wide methylation profiles can be reduced to compact panels suitable for experimental measurement ^15^. Restriction enzyme-based workflows further show that successful implementation depends on local sequence architecture and the availability of assay-compatible target sites ^16^. Integrating these components within a coordinate-resolved workflow enables methylation contrast and intersample variability to be considered together with application-relevant biological references and assay-compatible sequence features.

Here, we present a gene-centered analytical framework that integrates public DNA methylation profiles, phenotype metadata, pan-cancer and peripheral-blood reference layers, and matched gene-expression information where contextually applicable to prioritize methylation regions defined by explicit genomic start and end coordinates within specified biological contexts. Candidate regions are evaluated according to methylation contrast, intersample variability, compatibility with relevant biological backgrounds and local sequence features associated with downstream assay development. We applied the framework to colorectal cancer (CRC) and hepatocellular carcinoma (HCC), using cohort-specific tissue references during computational prioritization and disease-relevant tissue comparators during experimental evaluation. Selected regions were then evaluated experimentally with locus-specific Delta-HLD methylation assays against matched non-tumor colon tissue and cirrhotic liver tissue, respectively. The analytical development and panel-level implementation of Delta-HLD are reported in the companion study by Vaquer et al. ^17^. We also applied the framework to lung adenocarcinoma and lung squamous cell carcinoma to examine the effects of alternative tumor backgrounds and cohort composition on regional prioritization.

Together, these analyses establish a reproducible path from genome-scale methylation data to context-prioritized genomic intervals with explicit coordinates and assay-relevant sequence features by defining the reference states, background tissues and tumor contexts that shape candidate selection.

## Results

### A gene-centered framework delineates context-aware methylation regions

We developed a gene-centered analytical framework that integrates public DNA methylation profiles, phenotype metadata, CpG annotations, matched RNA-seq data and peripheral-blood methylation references into coordinate-resolved analytical features (Fig. 1a and Extended Data Fig. 1). To establish its operating logic, we applied the framework to TCGA-COAD (The Cancer Genome Atlas-Colon Adenocarcinoma) methylation profiles from 292 tumors (T) and 38 non-tumor (NT) samples classified by TCGA as solid tissue normal and used SEPTIN9 as a clinically established benchmark locus ^18^. Gene-centered visualization revealed heterogeneous methylation across the broader locus, with tumor and non-tumor profiles diverging within a restricted genomic interval (Fig. 1b and Extended Data Fig. 2a).

**Fig. 1.**
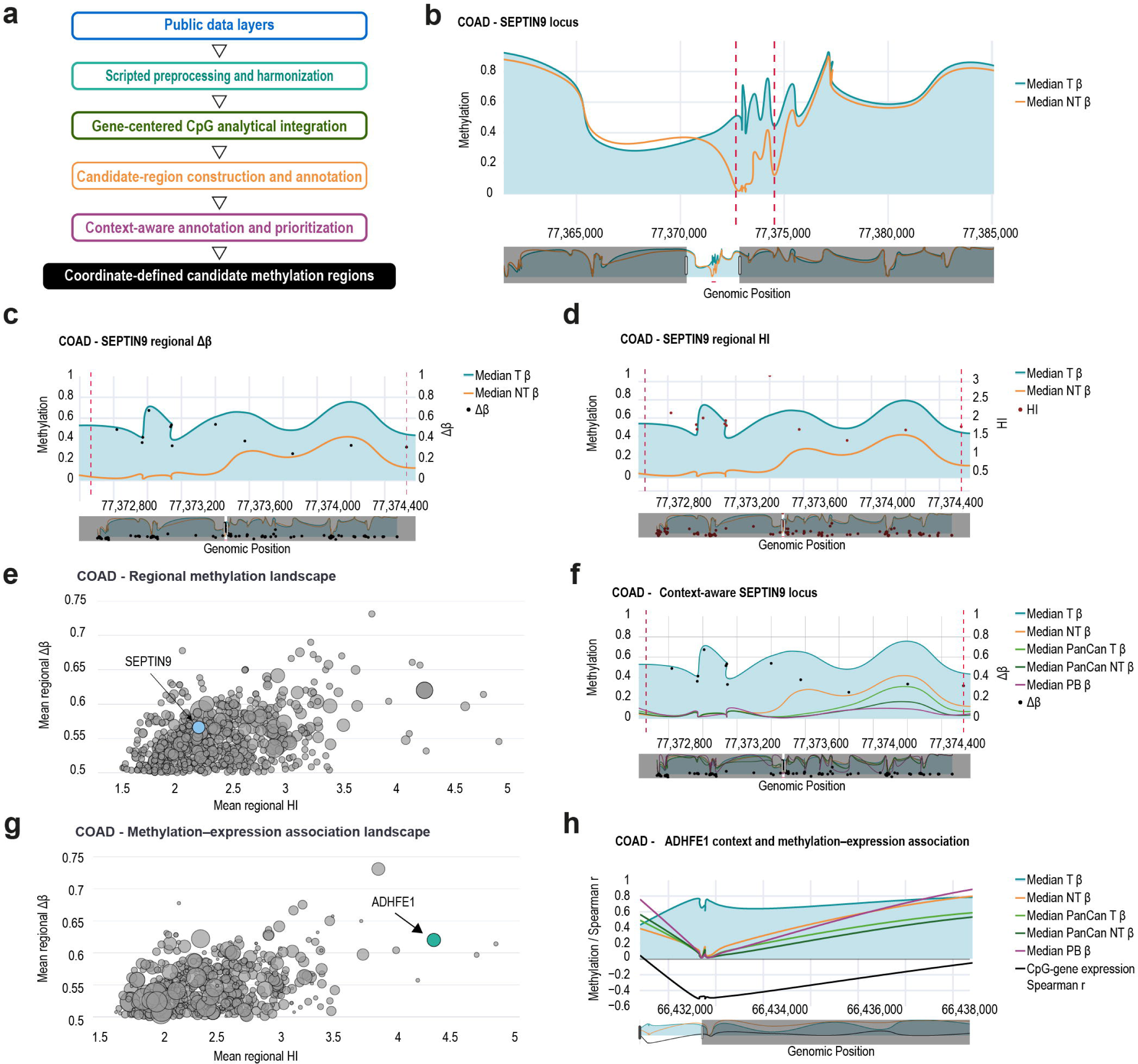
A gene-centered framework delineates context-aware methylation regions. **a,** Overview of the analytical framework, comprising public data layers, scripted preprocessing and harmonization, gene-centered CpG analytical integration, candidate region construction and annotation, and context-aware annotation and prioritization. **b,** Median methylation profiles across the SEPTIN9 locus in 292 TCGA-COAD tumors and 38 non-tumor samples classified by TCGA as solid tissue normal. Dashed lines indicate the interval examined in **c**, **d** and **f**. **c,** Regional SEPTIN9 profile showing median tumor and non-tumor β-values and CpG-level methylation differences (Δβ). Black points indicate CpG-level Δβ values. **d,** Regional SEPTIN9 profile showing CpG-level homogeneity index (HI) values. Red points indicate CpG-level HI values. The candidate region spans chr17:77,372,667-77,374,528 in GRCh38. **e,** Exploratory COAD regional methylation landscape obtained using Δβ ≥ 0.50 and HI ≥ 1.0. Each bubble represents a gene-centered region positioned by mean regional HI and mean regional Δβ; bubble size indicates the number of qualifying CpGs. SEPTIN9 is highlighted. **f,** Context-aware SEPTIN9 profile showing median β-values for COAD tumors, COAD non-tumor samples, 7,946 non-target pan-cancer tumors, 684 pan-cancer non-tumor samples and 139 peripheral blood samples, together with CpG-level Δβ values. COAD samples were excluded from the pan-cancer tumor reference. **g,** COAD methylation-expression association landscape. Regions are positioned by mean regional HI and mean regional Δβ; bubble size represents the strength of the inverse CpG-gene expression association. ADHFE1 is highlighted. **h,** ADHFE1 contextual methylation profile and CpG-gene expression Spearman correlation coefficients across TCGA-COAD tumors. The black line indicates the CpG-gene expression Spearman correlation coefficient. Lower tracks in **b-d**, **f** and **h** show the corresponding full-locus context. T, tumor; NT, non-tumor; PanCan, pan-cancer; PB, peripheral blood; Δβ, median tumor β minus median non-tumor β; HI, homogeneity index.

At CpG resolution, tumor-associated methylation was quantified using the median methylation difference between tumor and non-tumor samples, Δβ, together with a variability-adjusted contrast, the Homogeneity Index (HI), defined as |Δβ|/√(SD_T² + SD_NT²). Δβ captured the magnitude of the methylation shift. HI downweighted CpGs showing high intersample dispersion. Concordant increases in both metrics delineated a localized SEPTIN9 candidate methylation region supported by coordinated CpG-level evidence at chr17:77,372,667-77,374,528 (GRCh38) (Fig. 1c,d and Extended Data Fig. 2b,c). Extension of this analysis across the TCGA-COAD methylome under an exploratory configuration of Δβ ≥ 0.50 and HI ≥ 1.0 retained 1,030 CpGs distributed across 714 gene-centered regions. Positioning these regions by their mean Δβ and HI values provided an exploratory regional methylation landscape, with SEPTIN9 serving as an internal reference (Fig. 1e, Supplementary Table 1).

We next evaluated regional behavior across biological background layers. The framework integrated methylation summaries from the target tumor and non-tumor cohorts with 7,946 non-target pan-cancer tumors (PanCan T) across 32 cancer types and subtypes, 684 pan-cancer non-tumor samples (PanCan NT) across 23 organs, both derived from the GDC-TCGA Pan-Cancer HumanMethylation450K dataset, together with 139 peripheral-blood (PB) methylation profiles obtained from GEO series GSE270856 and GSE247869 (Fig. 1f and Extended Data Fig. 2d). These overlays enabled potentially confounding methylation outside the primary comparison between tumor and non-tumor tissue to be inspected directly. The COAD cohort was excluded from the pan-cancer tumor reference to prevent the same samples from contributing to both the target cohort and the background reference.

Matched RNA-seq data were incorporated as an orthogonal annotation by calculating Spearman correlations between CpG methylation and gene expression across tumor samples. The resulting methylation-expression association landscape highlighted regions in which tumor-associated hypermethylation was accompanied by an inverse association with expression, without making this association a universal requirement for regional retention (Fig. 1g and Supplementary Table 2). ADHFE1 illustrated this contextual integration by combining favorable regional Δβ and HI values with inverse methylation-expression associations across the evaluated interval (Fig. 1h and Extended Data Fig. 2e).

Together, these analyses establish the operating principle of the framework. Array CpGs provide coordinate-level evidence, adjacent signals are aggregated into candidate methylation regions, and biological background and expression layers support contextual interpretation before contextual candidate assessment and assay-aware sequence inspection.

### Context-aware filtering prioritizes coordinate-defined methylation regions in COAD

Having established the framework’s operating principle, we applied a multilayer configuration to prioritize methylation regions defined by explicit genomic coordinates across the TCGA-COAD methylome (Fig. 2a). The screening universe comprised 253,334 CpGs mapped to 21,486 genes and 58,860 gene-centered regions. For each CpG, the analytical layer integrated target tumor and non-tumor methylation, pan-cancer tumor and non-tumor methylation references, peripheral-blood methylation, Δβ, HI and the association between CpG methylation and gene expression.

**Fig. 2.**
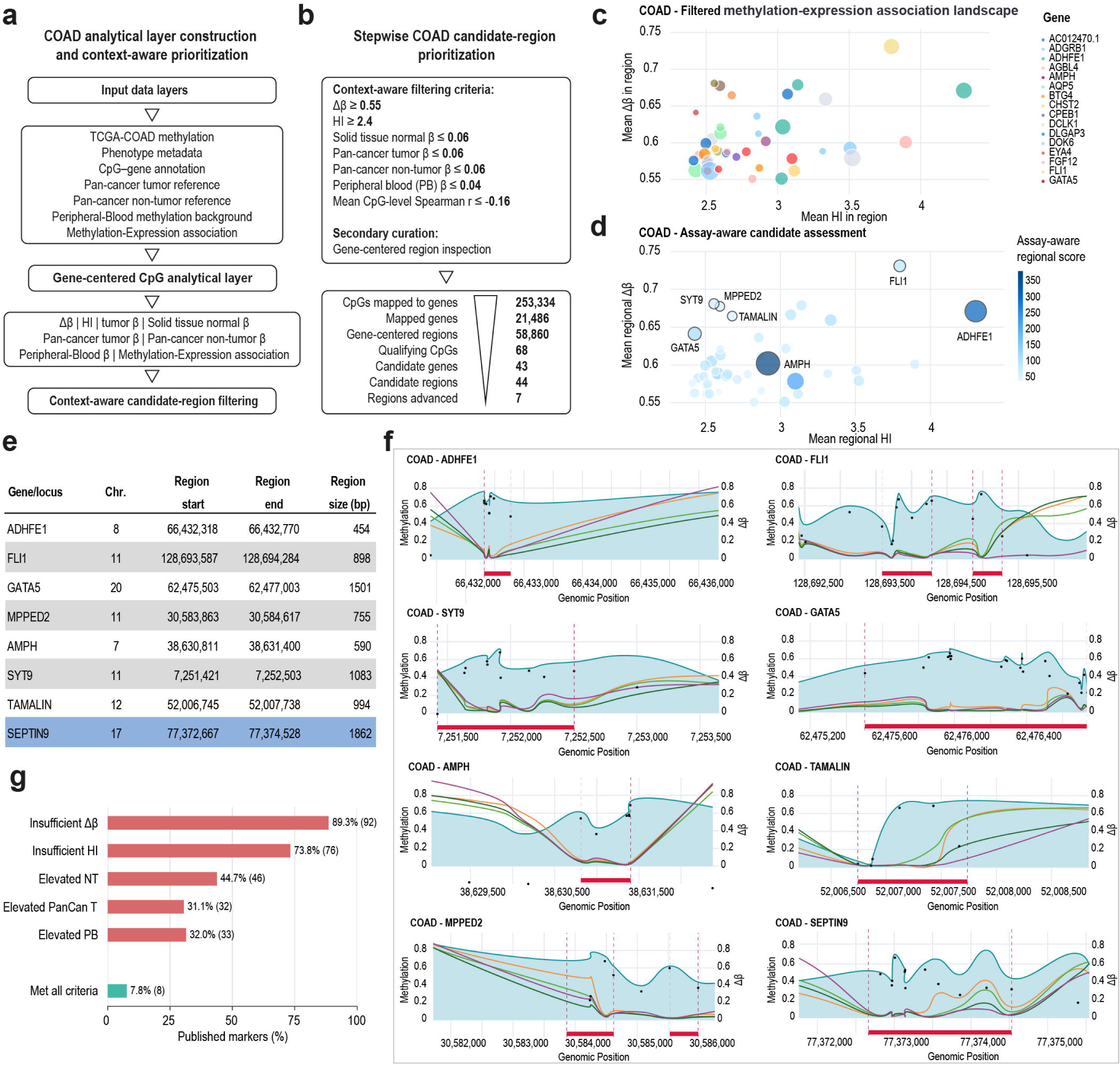
Context-aware prioritization of coordinate-defined methylation regions in TCGA-COAD. **a,** Construction of the TCGA-COAD analytical layer and context-aware candidate region prioritization. Input data comprise TCGA-COAD methylation and phenotype data, CpG-gene annotations, pan-cancer tumor and non-tumor reference layers, peripheral blood methylation profiles and CpG-gene expression associations. **b,** Stepwise filtering of 253,334 CpGs mapped to 21,486 genes and 58,860 gene-centered regions. Context-aware criteria retained 68 qualifying CpGs mapping to 43 genes and 44 candidate regions. Seven regions were subsequently advanced for sequence-level evaluation following multidimensional regional and sequence assessment. **c,** Filtered methylation-expression association landscape containing the 44 candidate regions. Each bubble is positioned according to mean regional HI and mean regional Δβ; bubble size represents the strength of the inverse CpG-gene expression association, and color denotes gene identity. Regions with identical or similar summary values may overlap in the static representation. **d,** Assay-aware assessment of the filtered candidate regions. Bubble size and color indicate an assay-aware regional score integrating regional CpG support, inverse methylation-expression association and sequence-derived features relevant to downstream HhaI-sensitive Delta-HLD implementation. The score was used as an auxiliary candidate annotation and not as a rank-order selection criterion. The seven regions advanced for sequence-level evaluation are labeled. **e,** Genomic coordinates and sizes of the seven regions advanced for sequence-level evaluation and the clinically established SEPTIN9 benchmark. **f,** Gene-centered methylation profiles of the selected loci and SEPTIN9, showing median β-values for COAD tumors, COAD non-tumor samples, non-target pan-cancer tumors, pan-cancer non-tumor samples and peripheral blood samples, together with CpG-level Δβ values. Red horizontal bars indicate coordinate-defined candidate regions. For FLI1, two candidate regions were identified; the solid red bar indicates the region advanced for sequence-level inspection. Lower tracks show the corresponding full-locus context. **g,** Frequency of criteria not met among 103 literature-reported CRC methylation markers. Bars show percentages, with the number of markers indicated in parentheses. Individual markers could fail more than one criterion; marker-level results are provided in Supplementary Data 1. T, tumor; NT, non-tumor; PanCan, pan-cancer; PB, peripheral blood; Δβ, median tumor β minus median non-tumor β; HI, homogeneity index.

Context-aware methylation filtering was first performed at CpG resolution using Δβ ≥ 0.55, HI ≥ 2.40, median solid tissue normal β ≤ 0.06, median pan-cancer tumor β ≤ 0.06, median pan-cancer non-tumor β ≤ 0.06 and median peripheral-blood β ≤ 0.04. CpGs passing these criteria were mapped to the predefined physical regions, and CpG-level methylation-expression associations were summarized within each region. Regions with a mean CpG-level Spearman coefficient ≤ −0.16 were subsequently retained as an additional COAD-specific prioritization step. The complete procedure retained 68 CpGs mapping to 43 genes and 44 candidate methylation regions (Fig. 2b).

The 44 candidate regions were positioned within a regional prioritization landscape defined by mean regional Δβ and HI, with methylation-expression association and assay-aware information providing additional dimensions for candidate assessment (Fig. 2c,d and Supplementary Tables 3 and 4). The assay-aware score integrated CpG support, inverse methylation-expression association and sequence-derived features relevant to downstream HhaI-sensitive Delta-HLD implementation and was used as an auxiliary regional annotation rather than as a rank-order selection criterion.

A subset of seven regions was advanced for downstream sequence evaluation following structured review of the regional prioritization landscape, providing a practical experimental sampling of complementary positions across the candidate space. ADHFE1 represented a region with particularly high HI, whereas FLI1 combined high Δβ and HI. TAMALIN, SYT9 and MPPED2 occupied a neighboring sector characterized by high Δβ, AMPH showed a high assay-aware score, and GATA5 combined high Δβ with comparatively lower HI, providing a contrasting regional profile within the landscape. These regions therefore sampled distinct positions within the prioritization space while retaining compatible biological background and sequence characteristics. SEPTIN9 was retained as the clinically established benchmark (Fig. 2d).

For each region advanced for sequence evaluation, the framework returned chromosome, start and end coordinates, and interval size (Fig. 2e). Gene-centered inspection localized the prioritized intervals within the broader methylation profiles and enabled their behavior across target and background layers to be reviewed before sequence-level analysis (Fig. 2f). A single gene-centered locus could contain more than one candidate interval, as illustrated by the two regions identified within FLI1, one of which was advanced for downstream sequence inspection. Across the selected loci, advancement to sequence-level evaluation reflected the combined consideration of regional Δβ and HI, CpG support, background compatibility, methylation-expression association, local methylation coherence and sequence suitability.

An audit of 103 published CRC methylation markers compiled from the literature (representative studies^18–22^; complete source list in Supplementary Data 1) showed that many did not satisfy the context-aware criteria applied here. Insufficient Δβ was the most frequent limitation, affecting 92 markers (89.3%), followed by insufficient HI in 76 (73.8%), elevated non-tumor tissue methylation in 46 (44.7%), elevated peripheral-blood methylation in 33 (32.0%) and elevated pan-cancer tumor methylation in 32 (31.1%). Only eight genes (7.8%) met all evaluated criteria. Individual markers could fail multiple criteria (Fig. 2g and Supplementary Data 1).

In combination, context-aware filtering reduced the TCGA-COAD methylome to 44 candidate methylation regions. Seven regions, together with the SEPTIN9 benchmark, were advanced for sequence-resolved evaluation and defined constrained genomic search spaces for subsequent identification of assay-compatible GCGC sites.

### Sequence-resolved methylation targets distinguish CRC from normal-adjacent tissue

Following the sequence review described above, the seven COAD regions advanced for experimental evaluation were examined at nucleotide resolution to identify experimentally addressable methylation targets. The framework retrieved the corresponding GRCh38 reference sequences and annotated array-manifest CpGs, total CpG content, GC fraction and GCGC motifs (Fig. 3a). GCGC motifs were included as assay-relevant sequence features because they define sites amenable to methylation-sensitive HhaI digestion, providing a direct route from prioritized regions to targeted methylation analysis in native DNA. The exported sequence view comprised the prioritized core interval flanked by 100-bp sequences on either side, enabling contextual inspection of manifest CpGs and GCGC motif distribution. At the ADHFE1 locus, the displayed sequence spanned chr8:66,432,217-66,432,870 and contained a prioritized core interval at chr8:66,432,317-66,432,770, with a GC fraction of 0.639 and nine GCGC motifs. Candidate GCGC sites were then nominated through structured visual review of the exported sequence context, considering motif position within the prioritized interval, local CpG density and surrounding sequence composition. Regional sequence views for the remaining prioritized COAD loci and the SEPTIN9 benchmark are shown in Supplementary Data 2.

**Fig. 3.**
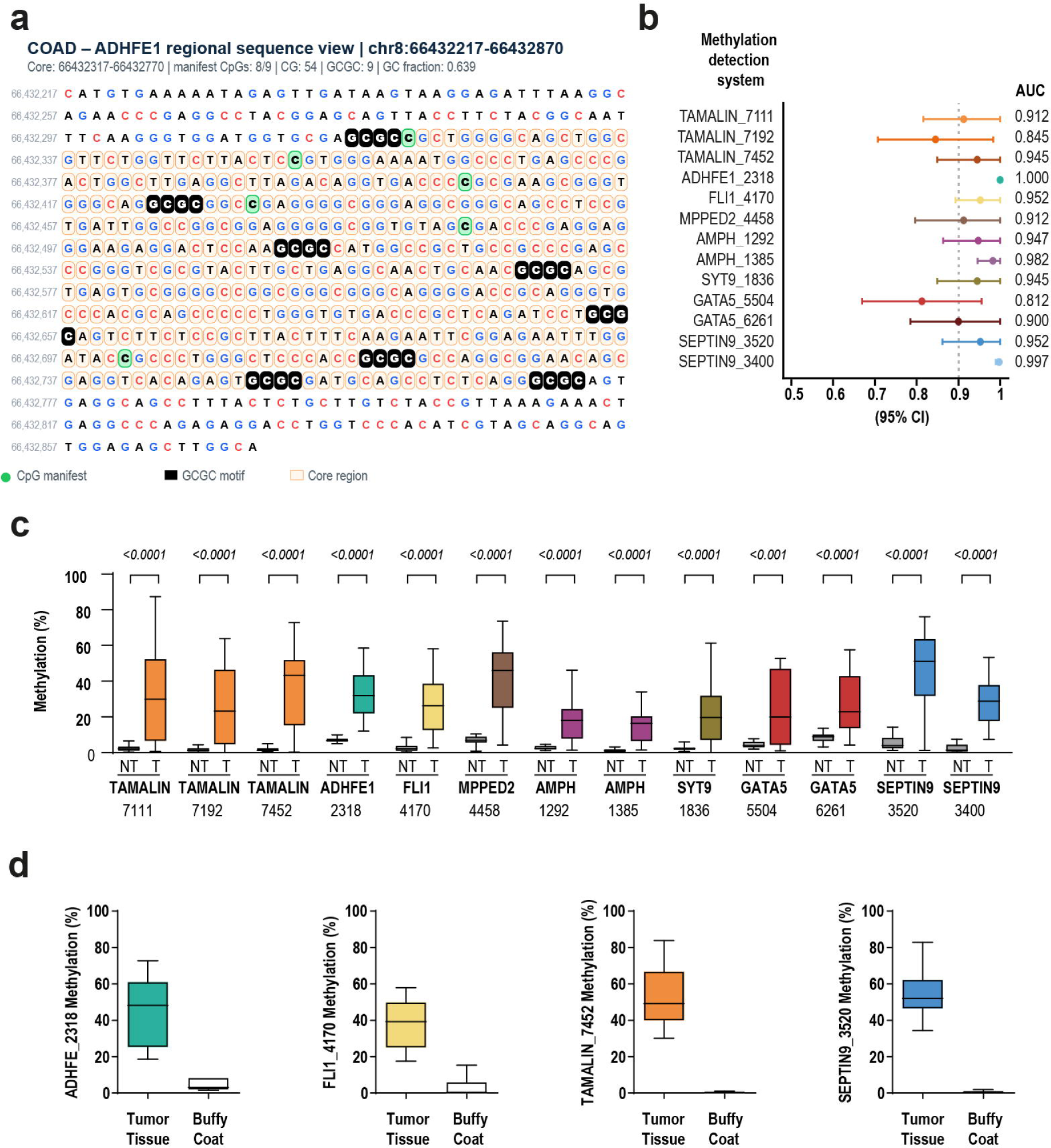
Sequence-resolved methylation targets derived from the seven regions advanced for sequence evaluation distinguish CRC tumors from normal-adjacent tissue. **a,** Regional sequence view exported from the framework for the ADHFE1 locus in TCGA-COAD. The displayed sequence spans chr8:66,432,217-66,432,870 in GRCh38 and includes the prioritized core interval at chr8:66,432,317-66,432,770 together with 100 bp of flanking sequence on each side. CpGs represented on the array manifest, GCGC motifs and the prioritized core region are indicated. **b,** Tissue discrimination performance of the 13 methylation detection systems derived from the seven prioritized regions and the SEPTIN9 benchmark, shown as AUC with 95% confidence intervals. The dashed line indicates an AUC of 0.90. **c,** Methylation levels measured by qPCR-based Delta-HLD in fresh-frozen CRC tumors and paired normal-adjacent tissues from 20 resections. All systems showed higher median methylation in tumor tissue than in the corresponding normal-adjacent tissue. P values are shown above each comparison. **d,** Methylation levels for four representative assays in tumor tissue and patient-derived buffy coat DNA. ADHFE1_2318, FLI1_4170, TAMALIN_7452 and SEPTIN9_3520 showed higher methylation in tumor tissue and low methylation in buffy coat DNA. NT, normal-adjacent tissue; T, tumor; AUC, area under the receiver operating characteristic curve.

Selected GCGC sites were subsequently used to design 13 locus-specific Delta-HLD methylation assays across the seven prioritized regions and the SEPTIN9 benchmark, with some regions supporting more than one assay. These assays were evaluated in fresh-frozen tumors and paired normal-adjacent tissues from 20 CRC resections. All 13 assays showed higher median methylation in tumor tissue than in the corresponding normal-adjacent tissue (Fig. 3b,c). AUC values for tissue discrimination ranged from 0.812 to 1.000, with a median of 0.945. Eleven of the 13 assays reached AUCs of at least 0.90 and eight reached AUCs of at least 0.945. ADHFE1_2318 achieved an AUC of 1.000, followed by SEPTIN9_3400 at 0.997 and AMPH_1385 at 0.982.

We additionally evaluated four representative assays in tumor tissue and buffy coat DNA obtained from patients with CRC. ADHFE1_2318, FLI1_4170, TAMALIN_7452 and SEPTIN9_3520 showed higher methylation in tumor tissue and low methylation in buffy coat DNA (Fig. 3d), consistent with the low hematopoietic background predicted by the peripheral-blood reference layer used during computational prioritization.

Together, regional sequence annotation, structured site selection and assay design connected COAD-derived evidence with locus-specific Delta-HLD methylation assays that discriminated CRC tumors from matched normal-adjacent tissues.

### Context-aware prioritization identifies HCC-associated methylation regions that distinguish HCC from cirrhotic liver

Having established the operating logic of the framework in COAD, we next extended it to HCC, where the intended clinical comparison was discrimination from cirrhotic liver. The analytical layer comprised HumanMethylation450K profiles from 375 LIHC tumors and 50 adjacent non-tumor liver samples from patients with HCC. These TCGA samples provided the non-neoplastic liver reference for computational prioritization; the resulting regions were subsequently evaluated directly against histologically confirmed cirrhotic liver in the independent cohort.

Context-aware filtering used Δβ ≥ 0.46, HI ≥ 1.5, median LIHC non-tumor liver β ≤ 0.05, median pan-cancer tumor β ≤ 0.07, median pan-cancer non-tumor β ≤ 0.06 and median peripheral-blood β ≤ 0.04. For LIHC, the prioritization configuration was restricted to methylation contrast, biological background and sequence features. Because the available methylation-expression layer represented covariation within HCC tumors rather than reference-state evidence directly matched to the intended HCC-versus-cirrhosis evaluation, it was not incorporated into candidate filtering or the assay-aware regional annotation (Fig. 4a). These criteria reduced the shared analytical universe of 253,334 CpGs, 21,486 genes and 58,860 gene-centered regions to 19 qualifying CpGs mapping to 15 genes and 16 candidate regions.

**Fig. 4.**
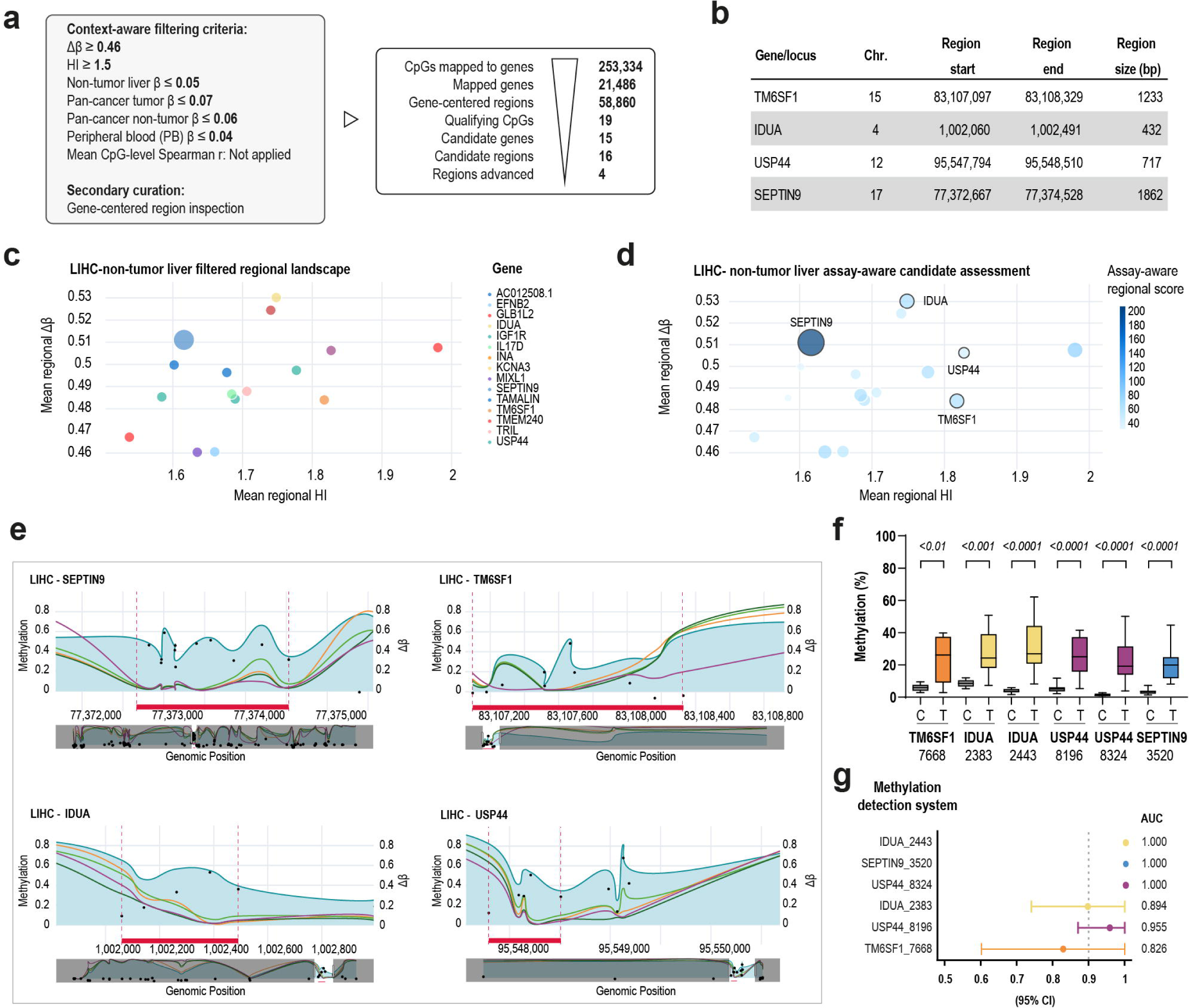
Context-aware prioritization identifies HCC-associated methylation regions that distinguish HCC from cirrhotic liver. **a,** Context-aware filtering of TCGA-LIHC methylation data using adjacent non-tumor liver from patients with HCC as the reference. Filtering of 253,334 CpGs mapped to 21,486 genes and 58,860 gene-centered regions retained 19 qualifying CpGs, 15 genes and 16 candidate regions, of which four were advanced for sequence-resolved evaluation following regional and sequence assessment. Methylation-expression information was treated as a context-dependent layer and was not incorporated into LIHC candidate filtering or regional scoring. **b,** Genomic coordinates and sizes of the four prioritized regions. **c,** Filtered regional landscape for LIHC versus adjacent non-tumor liver. Each bubble represents a candidate region positioned by mean regional HI and mean regional Δβ; bubble size indicates the number of qualifying CpGs and color denotes gene identity. **d,** Assay-aware regional prioritization of the 16 candidate regions. Bubble size and color indicate the regional score adapted for the LIHC comparison, which excluded the methylation-expression component. The score provided an auxiliary assay-aware annotation and was not used as a rank-order selection criterion. The four regions advanced for sequence-resolved evaluation are labeled. **e,** Gene-centered methylation profiles for SEPTIN9, TM6SF1, IDUA and USP44, showing median β-values for LIHC tumors, LIHC adjacent non-tumor liver, non-target pan-cancer tumors, pan-cancer non-tumor samples and peripheral blood samples, prioritized regions; lower tracks show the corresponding full-locus context. **f,** qPCR-based Delta-HLD methylation levels for six methylation detection systems in 11 HCC tumors and 12 cirrhotic liver specimens. All six systems showed higher median methylation in HCC tumors than in cirrhotic liver specimens. P values are shown above each comparison. **g,** Tissue discrimination AUCs with 95% confidence intervals for the six systems. The dashed line indicates an AUC of 0.90. C, cirrhotic liver; T, HCC tumor; PanCan, pan-cancer; PB, peripheral blood; Δβ, median LIHC tumor β minus median LIHC non-tumor liver β; HI, homogeneity index; AUC, area under the receiver operating characteristic curve; MSRE, methylation-sensitive restriction enzyme.

The filtered regions were positioned according to mean regional Δβ and HI, with an assay-aware regional score providing an additional annotation of CpG support and sequence characteristics relevant to downstream HhaI-sensitive Delta-HLD implementation (Fig. 4c,d and Supplementary Tables 5 and 6). The score was used as an additional candidate annotation and not as a rank-order selection criterion.

Candidates were evaluated within the regional prioritization landscape and through structured review of their local sequence context. Four regions mapping to TM6SF1, IDUA, USP44 and SEPTIN9 were advanced for sequence-resolved evaluation based on their combined regional, background and assay-relevant characteristics (Fig. 4b). SEPTIN9, previously used as a benchmark in the COAD analysis, also satisfied the criteria for the comparison between LIHC and its adjacent non-tumor liver reference. This finding shows that a methylation region observed across tumor contexts can remain informative when assessed against an HCC-associated non-tumor liver background.

Gene-centered profiles showed localized increases in LIHC tumor methylation relative to LIHC adjacent non-tumor liver, together with low peripheral-blood methylation and directly inspectable patterns across pan-cancer tumor and non-tumor references (Fig. 4e). Regional sequence views for TM6SF1, IDUA, USP44 and SEPTIN9, comprising each prioritized core interval and 100-bp flanking sequences, are shown in Supplementary Data 3. Sequence-resolved evaluation supported the design of six locus-specific Delta-HLD methylation assays, IDUA_2383, IDUA_2443, USP44_8196, USP44_8324, TM6SF1_7668 and SEPTIN9_3520.

The six assays were evaluated in an independently recruited patient cohort comprising 23 patient-derived liver specimens, including 11 HCC tumors and 12 cirrhotic liver control tissues. All six systems showed higher median methylation in HCC than in cirrhotic liver tissue (Fig. 4f). AUCs for liver tissue discrimination ranged from 0.826 to 1.000 (Fig. 4g). IDUA_2443, USP44_8324 and SEPTIN9_3520 achieved AUCs of 1.000, while USP44_8196, IDUA_2383 and TM6SF1_7668 achieved AUCs of 0.955, 0.894 and 0.826, respectively.

These results extend context-aware regional prioritization from COAD to HCC using an HCC-associated non-tumor liver reference. They further show that LIHC-derived regional evidence can guide the selection of locus-specific assays that distinguish HCC from cirrhotic liver tissue in an independent cohort.

### Histological subtype and cohort composition shape methylation region prioritization in lung cancer

We next evaluated how histological subtype within the same organ influences context-aware methylation region prioritization. LUAD (Lung Adenocarcinoma) and LUSC (Lung Squamous Cell Carcinoma) were configured as independent analytical contexts using HumanMethylation450K profiles from 455 LUAD tumors with 32 corresponding non-tumor samples and 365 LUSC tumors with 41 corresponding non-tumor samples, with the non-tumor specimens classified by TCGA as solid tissue normal. To place the lung subtype comparison within the range observed across the previously analyzed disease contexts, we first compared candidate region yields using a common Δβ ≥ 0.50 threshold, with the remaining filters left unrestricted. This configuration retained 714 candidate regions in COAD, 90 in LIHC, 32 in LUAD and 264 in LUSC, corresponding to 1.21%, 0.15%, 0.05% and 0.45% of the shared universe of 58,860 regions, respectively (Fig. 5a and Extended Data Fig. 3a,b and Supplementary Tables 7 and 8). Candidate region yield therefore varied substantially across analytical contexts and differed by 8.3-fold between LUAD and LUSC despite application of the same Δβ threshold. This analysis was designed to examine how subtype-specific biological context shapes methylation-region prioritization within the discovery framework.

**Fig. 5.**
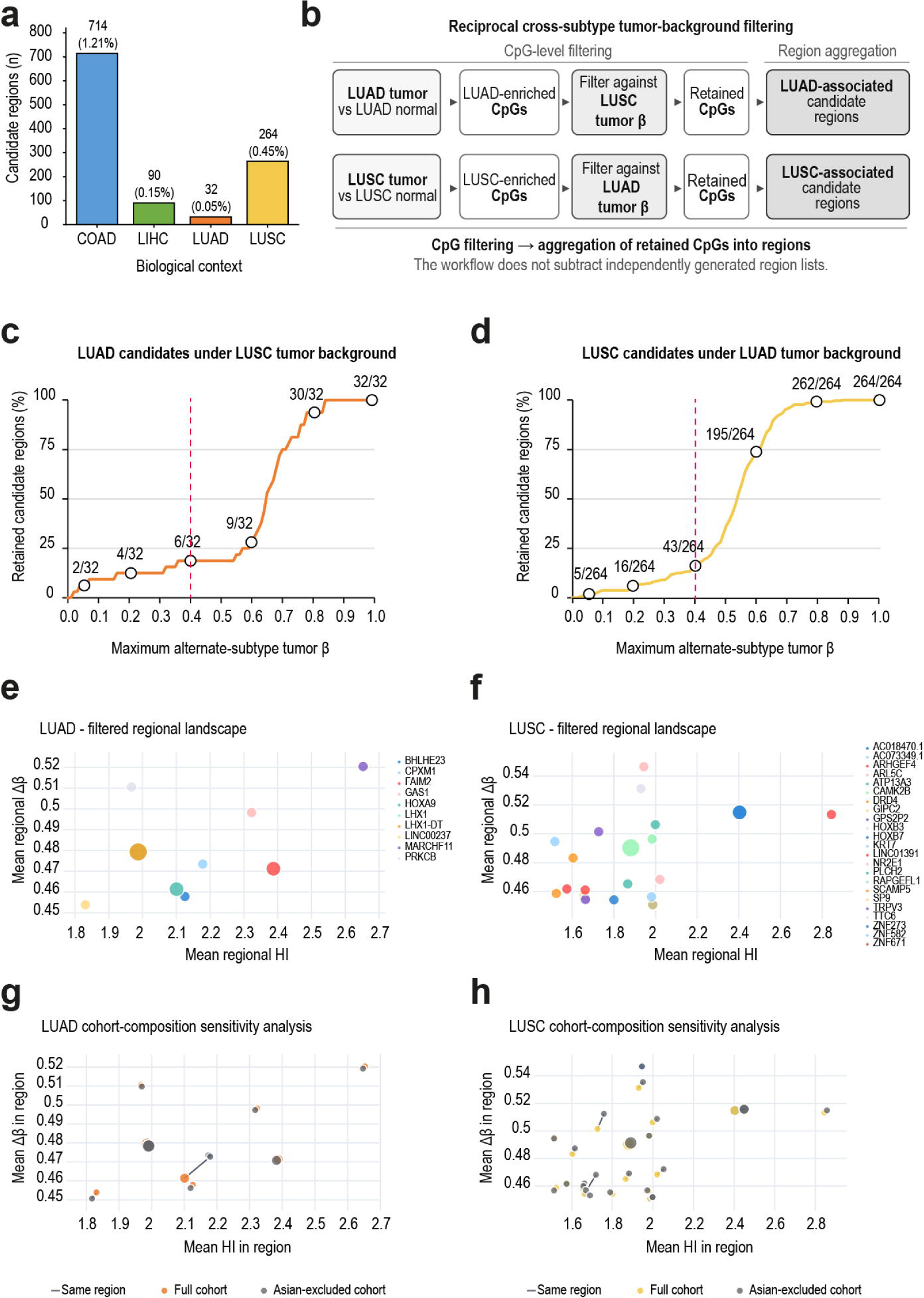
Reciprocal tumor background filtering and cohort composition shape lung methylation region prioritization. **a,** Candidate region yields across four biological contexts using a common Δβ ≥ 0.50 criterion, with the remaining context filters set to non-restrictive values. Percentages indicate the proportion of the shared universe of 58,860 gene-centered regions retained in each context. **b,** Reciprocal filtering against the alternate subtype tumor reference layer. CpGs enriched in LUAD relative to LUAD non-tumor samples classified by TCGA as solid tissue normal were filtered against the LUSC tumor reference layer, and LUSC-enriched CpGs were filtered reciprocally against the LUAD tumor reference layer. Filtering was performed at CpG resolution before aggregation of retained CpGs into regions. **c,d,** Percentage of initial LUAD (**c**) and LUSC (**d**) candidate regions retained across increasing maximum β-values permitted in the alternate subtype tumor reference layer, using the exploratory Δβ ≥ 0.50 candidate sets shown in **a**. Fractions indicate the number of retained regions relative to the initial candidate set. Dashed red lines indicate the maximum alternate subtype tumor β-value of 0.40 used in the intermediate configuration. **e,f,** Filtered regional landscapes obtained using the intermediate configuration for LUAD (**e**) and LUSC (**f**). This configuration used Δβ ≥ 0.45, incorporated the contextual background filters and permitted a maximum alternate subtype tumor β-value of 0.40; the complete criteria are shown in Extended Data Fig. 3c,d. Each bubble represents a candidate region positioned by mean regional HI and mean regional Δβ; bubble size indicates the number of qualifying CpGs and color denotes gene identity. **g,h,** Cohort composition sensitivity analyses for LUAD (**g**) and LUSC (**h**). Points show regional summaries calculated using the full cohort and after excluding samples from patients annotated as Asian according to the patient-level *demographic.race* field in the TCGA/GDC clinical metadata; samples with other or unknown race annotations were retained. Lines connect regions matched by genomic coordinates, and bubble size indicates CpG support. The LUAD analysis included 455 tumors and 32 non-tumor samples in the full cohort and 449 tumors and 32 non-tumor samples after exclusion. The corresponding LUSC cohorts included 365 tumors and 41 non-tumor samples and 358 tumors and 40 non-tumor samples, respectively. COAD, colon adenocarcinoma; LIHC, liver hepatocellular carcinoma; LUAD, lung adenocarcinoma; LUSC, lung squamous cell carcinoma; Δβ, median target tumor β minus median corresponding non-tumor β; HI, homogeneity index.

To incorporate histological subtype directly into candidate selection, we next used the alternate histological subtype as an explicit tumor reference layer (Fig. 5b). CpGs enriched in LUAD relative to LUAD non-tumor samples were evaluated against LUSC tumor methylation, whereas LUSC-enriched CpGs were evaluated reciprocally against LUAD tumor methylation. Filtering against the alternate subtype was applied at CpG resolution before the retained CpGs were aggregated into regions. This procedure generated subtype-associated candidate regions without requiring subtraction of independently derived regional lists.

Using the exploratory Δβ ≥ 0.50 candidate sets shown in Fig. 5a, candidate region retention was quantified across increasing maximum β-values permitted in the tumor reference layer from the alternate subtype (Fig. 5c,d). Retention was nonlinear and differed between histologies. At a stringent alternate subtype limit of β ≤ 0.06, 2 of 32 LUAD regions and 5 of 264 LUSC regions were retained. At the intermediate β ≤ 0.40 threshold, 6 LUAD and 43 LUSC regions remained, whereas β ≤ 1.00 effectively disabled the filter against the alternate subtype and retained all initial regions. These results showed that the initial Δβ ≥ 0.50 threshold became highly restrictive when combined with filtering against the alternate histology, particularly in LUAD.

We therefore relaxed the Δβ threshold to 0.45 and incorporated the contextual filters previously applied in the other tumor settings, while maintaining the alternate subtype tumor reference at β ≤ 0.40. The complete filtering configuration is shown in Extended Data Fig. 3c,d. This configuration retained 13 qualifying CpGs mapping to 10 genes and 10 candidate regions in LUAD and 24 CpGs mapping to 23 genes and 23 candidate regions in LUSC (Fig. 5e,f and Supplementary Tables 9 and 10).

We then increased the stringency of the subtype comparison by reducing the maximum β-value permitted in the alternate tumor reference layer from 0.40 to 0.06, while retaining the remaining contextual filters. This configuration retained 9 qualifying CpGs mapping to 6 genes and 6 candidate regions in LUAD, and 7 qualifying CpGs mapping to 5 genes and 5 candidate regions in LUSC. The more stringent configurations and corresponding regional landscapes are shown in Extended Data Fig. 3e,f and Supplementary Tables 11 and 12. These outputs represent regions retained under stringent filtering against the alternate subtype and were not experimentally evaluated in the present study.

Finally, we examined whether candidate region prioritization was sensitive to the composition of the discovery cohort. The intermediate configuration was applied using the patient-level *demographic.race* field provided in the TCGA/GDC clinical metadata. Samples from patients annotated as Asian were excluded, while samples with other or unknown race annotations were retained. This reduced the LUAD cohort from 455 to 449 tumors without changing the 32 non-tumor samples, and reduced the LUSC cohort from 365 to 358 tumors and from 41 to 40 non-tumor samples. LUAD retained 10 candidate regions under both cohort definitions, although coordinate-matched regions shifted in mean regional Δβ, HI or CpG support (Fig. 5g and Supplementary Table 13). In LUSC, the retained set changed from 23 to 24 regions, accompanied by shifts in several shared intervals (Fig. 5h). Additional changes across the broader LUSC regional landscape are shown in Extended Data Fig. 3g, whereas coordinate-matched shifts are summarized in Extended Data Fig. 3h and Supplementary Table 14. This exclusion-based sensitivity analysis indicated that modest changes in cohort composition can alter regional summary metrics and candidate retention.

Together, these results extend context-aware prioritization to histological subtypes within the same organ and show that both the alternate subtype tumor reference layer and the composition of the discovery cohort can influence methylation region prioritization.

## Discussion

A DNA methylation signal does not carry fixed biomarker value in isolation, just as a coordinate becomes informative only within a map. Our results show that candidate retention and regional metrics depend on the biological reference states, background tissues and tumor contexts used during discovery. Integrating methylation contrast, intersample variability, pan-cancer and peripheral-blood references, matched gene expression and assay-relevant sequence features reduced genome-scale profiles to coordinate-defined regions. Biological context therefore guides candidate selection from the beginning of discovery.

The intended specimen type provides one example of application-specific biological context. In colorectal cancer, stool-based testing samples human genomic DNA through an anatomically restricted route, and multitarget assays incorporating methylated DNA markers have demonstrated detection of colorectal cancer and advanced precancerous lesions ^23,24^. Plasma presents a broader biological background because cell-free DNA originates from multiple tissues and the tumor-derived fraction is variable. Blood-based implementation may therefore require evaluation against hematopoietic and non-target tissue signals, together with other malignancies that may release overlapping methylation patterns ^5^. A prospective plasma assay also showed a distinct performance profile across colorectal cancer and advanced precancerous lesions ^8^, illustrating that biomarker performance can vary across biological states within the same intended application. Together, these examples indicate that specimen type and disease state should inform the biological reference backgrounds considered during biomarker discovery.

Regional aggregation added a further dimension to contextual prioritization. Cancer-associated methylation changes commonly span contiguous CpGs, and region-based methods can recover these events across array and bisulfite sequencing data ^25^. In our framework, Δβ quantified the methylation shift, whereas HI expressed that shift relative to the combined intersample dispersion of the tumor and reference cohorts. Their joint representation generated a context-specific landscape in which regions were positioned according to contrast and consistency. HI functions as an operational prioritization metric that integrates methylation contrast with intersample dispersion. Mapping qualifying CpGs to predefined gene-centered regions identifies candidate intervals, narrowing the sequence search space before experimental testing establishes individual site performance.

The analytical layers contributed complementary evidence, although their relevance depended on the biological context. Pan-cancer and peripheral-blood references exposed potentially confounding signals. In COAD, matched gene expression provided an orthogonal annotation for regions in which increased methylation was associated with reduced transcription^13^. Sequence analysis then identified GCGC motifs compatible with HhaI digestion. Restriction enzyme-based assays also depend on suitable recognition sites and assay-compatible sequences^16^. The experiments in patient-derived tissues showed that array-derived regional evidence can guide the design of locus-specific Delta-HLD methylation assays in independently recruited cohorts from a distinct clinical setting using a different analytical modality. The analytical development and panel-level implementation of Delta-HLD are described in the companion study by Vaquer et al. ^17^.

The experimentally evaluated regions remained discriminative in independently recruited tissue cohorts from a geographically and clinically distinct setting, showing that the underlying methylation differences were detectable beyond the TCGA discovery population. Their regional metrics and relative positioning within the prioritization landscape remain conditional on cohort composition. Recent tumor methylome studies have identified ancestry-associated DNA methylation differences that were validated across independent cohorts and remained evident across tumor-purity strata ^26^, and pan-cancer analyses have identified ancestry-associated methylation differences within TCGA cohorts ^27^. Much of the genetic control of DNA methylation is nevertheless shared across European and East Asian populations ^28^. Our lung cancer sensitivity analysis specifically assessed the sensitivity of regional prioritization to cohort composition. Excluding a small subgroup defined by self-reported race altered regional metrics and candidate retention without constituting a formal population-level comparison. Discovery in larger locally recruited cohorts characterized for genetic ancestry may preserve their relative positioning, modify it or reveal additional candidates.

SEPTIN9 further illustrates the application dependence of methylation biomarkers. Its clinical development established it as a plasma benchmark for colorectal cancer detection ^18^, whereas in our study the same locus satisfied the LIHC computational criteria and subsequently discriminated HCC from cirrhotic liver in the independent cohort. Prospective studies have also shown that circulating methylated SEPT9 contributes to hepatocellular carcinoma detection in patients with cirrhosis ^29,30^. This behavior is consistent with pan-cancer analyses identifying conserved and cancer-context-specific methylation signatures ^6,31^. The audit of published colorectal cancer biomarkers showed the same principle, as many candidates did not satisfy the contrast, dispersion or background constraints used here. Independent discovery studies have likewise incorporated normal tissue, blood controls and non-target malignancies when selecting markers for defined diagnostic contexts ^9,32^. Recent computational approaches further illustrate how analytical design can shape methylation candidate selection. Tang et al. addressed candidate reliability by accounting for measurement noise, individual characteristics and potentially causal disease associations ^14^, whereas Downs et al. showed that the formulation used to quantify methylation change alters the CpGs and biological processes prioritized from cancer methylomes ^33^. The present framework complements these approaches by carrying application-specific reference states and background layers through regional aggregation, contextual filtering and sequence-aware target selection. Published performance is shaped by the specimen types, reference backgrounds and analytical conditions used during development, and context-aware reassessment can inform the evaluation of candidates in a new application.

Several features define the current scope of the framework. Candidate retention and regional prioritization depend on public TCGA profiles, clinical metadata and preprocessing decisions. HumanMethylation450K interrogates a predefined subset of CpGs, and the gene-centered design excludes potentially informative intergenic regions. Pan-cancer medians provide interpretable references but compress heterogeneity within cancer types, tissues and cell populations. The thresholds remain exploratory and require prospective locking and validation. Peripheral-blood profiles served as computational references, and four CRC assays were additionally evaluated in buffy-coat DNA. The lung cancer regions remain experimentally unvalidated, and the race-based exclusion analysis supports conclusions only about cohort sensitivity. Within these boundaries, the framework provides a reproducible route from genome-scale methylation profiles to context-prioritized genomic intervals with explicit coordinates and assay-relevant sequence features. Experimental evaluation was restricted to selected subsets of the retained candidate regions, and the performance of the remaining computational candidates was not assessed. The conversion of selected intervals into locus-specific Delta-HLD methylation assays and their evaluation in patient-derived tissues establishes a direct link between context-aware computational prioritization and experimental measurement.

## Materials and Methods

### Data sources and integration

Public DNA methylation, gene-expression and phenotype data were obtained from the UCSC Xena/GDC Pan-Cancer resources for TCGA-COAD, TCGA-LIHC, TCGA-LUAD and TCGA-LUSC. Peripheral-blood methylation reference profiles were obtained from GEO series GSE247869 and GSE270856. Datasets were curated and harmonized to the GRCh38 genome assembly, integrating cohort-specific tumor and non-tumor profiles, non-target pan-cancer reference layers, peripheral-blood methylation and matched gene-expression information where applicable.

### Regional biomarker prioritization

CpG methylation profiles were organized within a gene-centered framework and summarized as coordinate-defined genomic regions. Candidate assessment integrated tumor-associated methylation contrast, intersample variability, methylation in application-relevant biological backgrounds and local sequence characteristics. Methylation-expression associations were incorporated when relevant to the analytical context. Regions were evaluated through a multidimensional assessment of regional methylation evidence and assay-compatible sequence features. Assay-aware scores served as auxiliary annotations during candidate review. Selected regions underwent sequence-level inspection to identify targets suitable for downstream experimental evaluation.

For lung cancer, LUAD and LUSC were evaluated using reciprocal filtering against the alternate histological subtype. Sensitivity to discovery-cohort composition was examined by repeating the analyses after excluding samples from patients annotated as Asian in TCGA/GDC clinical metadata, while retaining samples with other or unknown annotations and holding the analytical criteria constant.

### Clinical validation cohorts

Experimental evaluation used tissue specimens from the observational case-control studies CASCADE (NCT07148297) and HIDE (NCT07148310) in Mendoza, Argentina. Protocols were approved by the Ethics Committees of Hospital Español de Mendoza (Act No. 38/2022) and Hospital Central de Mendoza (Act No. 01/2023), respectively. All participants provided written informed consent, and the studies were conducted in accordance with the Declaration of Helsinki.

The CRC validation cohort comprised 20 resections contributing paired tumor and non-neoplastic colorectal tissue, with patient-derived buffy-coat DNA used for selected comparisons. The HCC validation set comprised 23 liver specimens from 16 individuals, including 11 HCC and 12 cirrhotic liver specimens. All specimens underwent histopathological evaluation.

### Methylation analysis

Genomic DNA was isolated from tissue and buffy-coat specimens using commercial extraction kits appropriate to each specimen type. Selected targets were evaluated using locus-specific Delta-HLD assays, a native-DNA methylation-sensitive enzymatic workflow coupled to quantitative PCR. Methylation levels were estimated from matched enzyme-treated and no-enzyme reactions, with technical replicate measurements combined for analysis. The analytical development and panel-level implementation of Delta-HLD are described in the companion study by Vaquer et al. 17.

### Statistical analysis

Statistical analyses were performed using GraphPad Prism. Patient-derived specimens were treated as the experimental units. CRC tumor and matched non-tumor tissue or buffy-coat measurements were compared using two-sided Wilcoxon matched-pairs signed-rank tests. HCC and cirrhotic liver measurements were compared using two-sided Mann-Whitney U tests. Discrimination was assessed by receiver operating characteristic analysis and summarized as area under the curve with 95% confidence intervals. Data are presented as median with interquartile range unless otherwise stated, and two-sided P values <0.05 were considered statistically significant.

## Data Availability

Publicly available DNA methylation, gene-expression and phenotype data analyzed in this study were obtained from the UCSC Xena/GDC Pan-Cancer (PANCAN) cohort, GDC Data Release 18.0 (released 8 July 2019; https://api.gdc.cancer.gov/data/), including the TCGA-COAD, TCGA-LIHC, TCGA-LUAD and TCGA-LUSC cohorts. Peripheral-blood methylation data are publicly available through the NCBI Gene Expression Omnibus under accession numbers GSE247869 and GSE270856. Processed data underlying the computational results reported in the manuscript, including the analytical values used to generate the figures and supplementary analyses, are provided as Supplementary Tables associated with this article. De-identified data supporting the colorectal and hepatocellular carcinoma tissue-validation analyses are available from the corresponding author upon reasonable request, subject to applicable ethical and data-protection requirements.

## Code Availability

The computational code underlying the analyses reported in this manuscript is maintained in a private, version-controlled repository. It is not publicly available due to intellectual-property considerations related to ongoing patent-protection and commercial-development activities. A fixed version of the code used to generate the reported results will be made available confidentially to editors and reviewers during peer review. Additional access can be requested from the corresponding author.

## Acknowledgements

We thank the participants in the CASCADE and HIDE studies and the clinical, surgical, pathology and laboratory teams involved in participant recruitment, specimen collection and processing, and clinical and histopathological characterization. We acknowledge the investigators and participants who generated and shared the TCGA and Gene Expression Omnibus datasets analyzed in this study. We also thank Fundación Escuela de Medicina Nuclear (FUESMEN) for providing access to laboratory facilities used during the experimental evaluation.

## Author contributions

E.M.C. conceived and supervised the study. A.R.A., C.G.S. and J.S.A. developed and applied the computational framework and performed the computational analyses. C.C.V., P.A.W., V.B. and R.D.M. contributed to experimental validation and data analysis. G.S., A.C., M.C., W.R.M., G.A.V., R.P.R., R.N.M., J.D.R., P.V. and L.S. contributed to participant recruitment, specimen collection and clinical or histopathological characterization. A.L. contributed to project coordination and resources. A.R.A., C.G.S., J.S.A. and E.M.C. drafted the manuscript. All authors contributed to manuscript review and approved the final version.

---

**Extended Data Fig. 1. Detailed architecture of the gene-centered analytical framework.** Public methylation, phenotype, RNA sequencing, peripheral blood and reference genome data are processed through scripted curation and harmonization workflows. Cohort methylation summaries, CpG-gene mapping, methylation-expression associations and peripheral blood reference features are integrated into a gene-centered CpG analytical layer. Adjacent qualifying CpGs are aggregated into regions with explicit genomic start and end coordinates and annotated with local sequence features, including CpG content and GCGC motifs. Context-aware filtering defines candidate regions, which are subsequently evaluated using regional landscapes, assay-aware annotation and locus-level sequence inspection to support selection for downstream experimental evaluation. PB, peripheral blood; HI, homogeneity index.

**Extended Data Fig. 2. Full-locus contextual profiles of SEPTIN9 and ADHFE1 in TCGA-COAD. a,** Full-locus SEPTIN9 methylation profile showing median β-values for TCGA-COAD tumors and non-tumor samples classified by TCGA as solid tissue normal. Dashed lines mark the candidate interval examined in Fig. 1b-f. **b,** Full-locus SEPTIN9 profile with CpG-level Δβ values, shown as black points. **c,** Full-locus SEPTIN9 profile with CpG-level HI values, shown as red points. **d,** Full-locus SEPTIN9 profile showing median β-values for COAD tumors, COAD non-tumor samples, non-target pan-cancer tumors, pan-cancer non-tumor samples and peripheral blood samples, together with CpG-level Δβ values. **e,** Full-locus ADHFE1 contextual methylation profile and CpG-gene expression Spearman correlation coefficients across TCGA-COAD tumors. The black line indicates the CpG-gene expression Spearman correlation coefficient. Lower tracks show the corresponding full-locus overview. T, tumor; NT, non-tumor; PanCan, pan-cancer; PB, peripheral blood; Δβ, median tumor β minus median non-tumor β; HI, homogeneity index.

**Extended Data Fig. 3. Exploratory, intermediate and stringent lung methylation region landscapes and LUSC cohort composition sensitivity. a,b,** Exploratory regional methylation landscapes for LUAD (**a**) and LUSC (**b**) obtained using Δβ ≥ 0.50, with the remaining context filters set to non-restrictive values. Each bubble represents a gene-centered candidate region positioned by mean regional HI and mean regional Δβ; bubble size indicates the number of qualifying CpGs and color denotes gene identity. **c,d,** Intermediate contextual filtering criteria and candidate attrition for LUAD (**c**) and LUSC (**d**). The configuration used Δβ ≥ 0.45, HI ≥ 1.5, a maximum alternate subtype tumor β-value ≤ 0.40 and the additional contextual filters shown in each panel. Filtering of 253,334 CpGs mapped to 21,486 genes and 58,860 gene-centered regions retained 13 qualifying CpGs mapping to 10 genes and 10 regions in LUAD, and 24 qualifying CpGs mapping to 23 genes and 23 regions in LUSC. **e,f,** Regional methylation landscapes obtained after stringent filtering against the alternate tumor subtype, using a maximum alternate subtype tumor β of 0.06 while retaining the remaining contextual criteria. The LUAD configuration retained 9 qualifying CpGs mapping to 6 genes and 6 candidate regions, whereas the LUSC configuration retained 7 qualifying CpGs mapping to 5 genes and 5 candidate regions. Bubble size represents CpG support and color denotes gene identity. **g,** LUSC cohort composition sensitivity analysis across the broader regional landscape. Regional summaries calculated using the full cohort are overlaid with those obtained after excluding samples from patients annotated as Asian according to the patient-level *demographic race* field in the TCGA/GDC clinical metadata; samples with other or unknown race annotations were retained. Lines connect regions matched by genomic coordinates. **h,** Coordinate-matched regional displacements between the two LUSC cohort definitions, shown without the corresponding bubbles to emphasize the magnitude and direction of regional shifts. The full LUSC cohort comprised 365 tumors and 41 non-tumor samples, and the cohort after exclusion comprised 358 tumors and 40 non-tumor samples. LUAD, lung adenocarcinoma; LUSC, lung squamous cell carcinoma; Δβ, median target tumor β minus median corresponding non-tumor β; HI, homogeneity index; PB, peripheral blood.

## References

1. Loyfer, N. et al. A DNA methylation atlas of normal human cell types. Nature 613, 355–364 (2023).

2. Zhu, T. et al. A pan-tissue DNA methylation atlas enables in silico decomposition of human tissue methylomes at cell-type resolution. Nat. Methods 19, 296–306 (2022).

3. Heeke, S. et al. Tumor- and circulating-free DNA methylation identifies clinically relevant small cell lung cancer subtypes. Cancer Cell 42, 225–237.e5 (2024).

4. Bie, F. et al. Multimodal analysis of cell-free DNA whole-methylome sequencing for cancer detection and localization. Nat. Commun. 14, 6042 (2023).

5. Conway, A.-M. et al. A cfDNA methylation-based tissue-of-origin classifier for cancers of unknown primary. Nat. Commun. 15, 3292 (2024).

6. Zeng, Y. et al. A pan-cancer compendium of 1,294 plasma cell-free DNA methylomes and fragmentomes enabling multicancer detection. Nat. Cancer 7, 384–398 (2026).

7. Mattox, A. K. et al. The origin of highly elevated cell-free DNA in healthy individuals and patients with pancreatic, colorectal, lung, or ovarian cancer. Cancer Discov. 13, 2166–2179 (2023).

8. Chung, D. C. et al. A cell-free DNA blood-based test for colorectal cancer screening. N. Engl. J. Med. 390, 973–983 (2024).

9. Cheishvili, D. et al. A high-throughput test enables specific detection of hepatocellular carcinoma. Nat. Commun. 14, 3306 (2023).

10. Guo, D. Z. et al. Early detection and prognosis evaluation for hepatocellular carcinoma by circulating tumour DNA methylation: a multicentre cohort study. Clin. Transl. Med. 14, e1652 (2024).

11. Yuan, D. et al. crossNN is an explainable framework for cross-platform DNA methylation-based classification of tumors. Nat. Cancer 6, 1283–1294 (2025).

12. Zhu, Z. et al. MethMarkerDB: a comprehensive cancer DNA methylation biomarker database. Nucleic Acids Res. 52, D1380–D1392 (2024).

13. Sakellaropoulos, T. et al. MethNet: a robust approach to identify regulatory hubs and their distal targets from cancer data. Nat. Commun. 15, 6027 (2024).

14. Tang, X. et al. Causality-driven candidate identification for reliable DNA methylation biomarker discovery. Nat. Commun. 16, 680 (2025).

15. Sun, M. et al. Tissue of origin prediction for cancer of unknown primary using a targeted methylation sequencing panel. Clin. Epigenetics 16, 25 (2024).

16. Vandenhoeck, J. et al. IMPRESS: Improved methylation profiling using restriction enzymes and smMIP sequencing, combined with a new biomarker panel, creating a multi-cancer detection assay. Br. J. Cancer 131, 1224–1236 (2024).

17. Vaquer, C. C. et al. Panel-level multilocus methylation quantification in native cell-free DNA by PCR-compatible sequential enzymatic processing. Preprint at medRxiv 10.64898/2026.06.11.26354897 (2026).

18. Church, T. R. et al. Prospective evaluation of methylated SEPT9 in plasma for detection of asymptomatic colorectal cancer. Gut 63, 317–325 (2014).

19. Lofton-Day, C. et al. DNA methylation biomarkers for blood-based colorectal cancer screening. Clin. Chem. 54, 414–423 (2008).

20. Lind, G. E. et al. Identification of an epigenetic biomarker panel with high sensitivity and specificity for colorectal cancer and adenomas. Mol. Cancer 10, 85 (2011).

21. Barták, B. K. et al. Colorectal adenoma and cancer detection based on altered methylation pattern of SFRP1, SFRP2, SDC2, and PRIMA1 in plasma samples. Epigenetics 12, 751–763 (2017).

22. Petit, J. et al. Evaluation of epigenetic methylation biomarkers for the detection of colorectal cancer using droplet digital PCR. Sci. Rep. 13, 8883 (2023).

23. Imperiale, T. F. et al. Multitarget stool DNA testing for colorectal-cancer screening. N. Engl. J. Med. 370, 1287–1297 (2014).

24. Imperiale, T. F. et al. Next-generation multitarget stool DNA test for colorectal cancer screening. N. Engl. J. Med. 390, 984–993 (2024).

25. Benelli, M. et al. Charting differentially methylated regions in cancer with Rocker-meth. Commun. Biol. 4, 1249 (2021).

26. Craddock, J. et al. Methylation reprogramming associated with aggressive prostate cancer and ancestral disparities. Mol. Syst. Biol. 21, 1676–1701 (2025).

27. Lee, K. K. et al. Association of genetic ancestry and molecular signatures with cancer survival disparities: a pan-cancer analysis. Cancer Res. 82, 1222–1233 (2022).

28. Hatton, A. A. et al. Genetic control of DNA methylation is largely shared across European and East Asian populations. Nat. Commun. 15, 2713 (2024).

29. Oussalah, A. et al. Circulating methylated SEPT9 for detection of hepatocellular carcinoma in cirrhosis. JAMA Oncol. 10.1001/jamaoncol.2026.2157 (2026).

30. Zheng, K. et al. Methylated SEPT9 combined with AFP and PIVKA-II is effective for the detection of HCC in high-risk population. BMC Gastroenterol. 23, 260 (2023).

31. Qin, Q. et al. Conserved methylation signatures associate with the tumor immune microenvironment and immunotherapy response. Genome Med. 16, 47 (2024).

32. Li, D. et al. Discovery and validation of tissue-specific DNA methylation as noninvasive diagnostic markers for colorectal cancer. Clin. Epigenetics 14, 102 (2022).

33. Downs, B. M. et al. An unbiased approach to measure aberrant DNA methylation alterations. Nat. Commun. 17, 4522 (2026).

