## Supplementary figures and images for "A context-aware analytical framework for DNA methylation biomarker discovery"

### Extended Data Fig 1

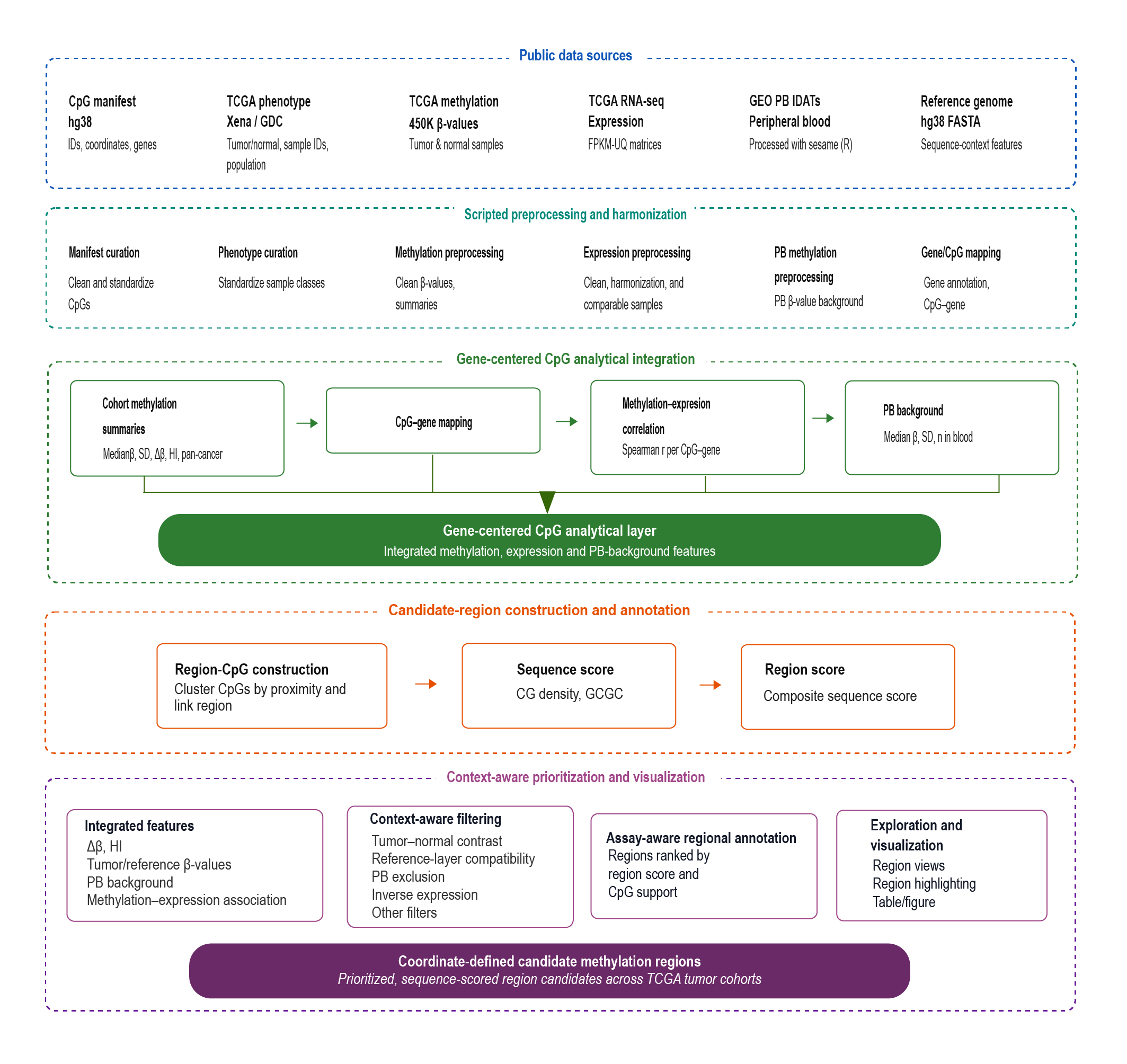

### Extended Data Fig 2

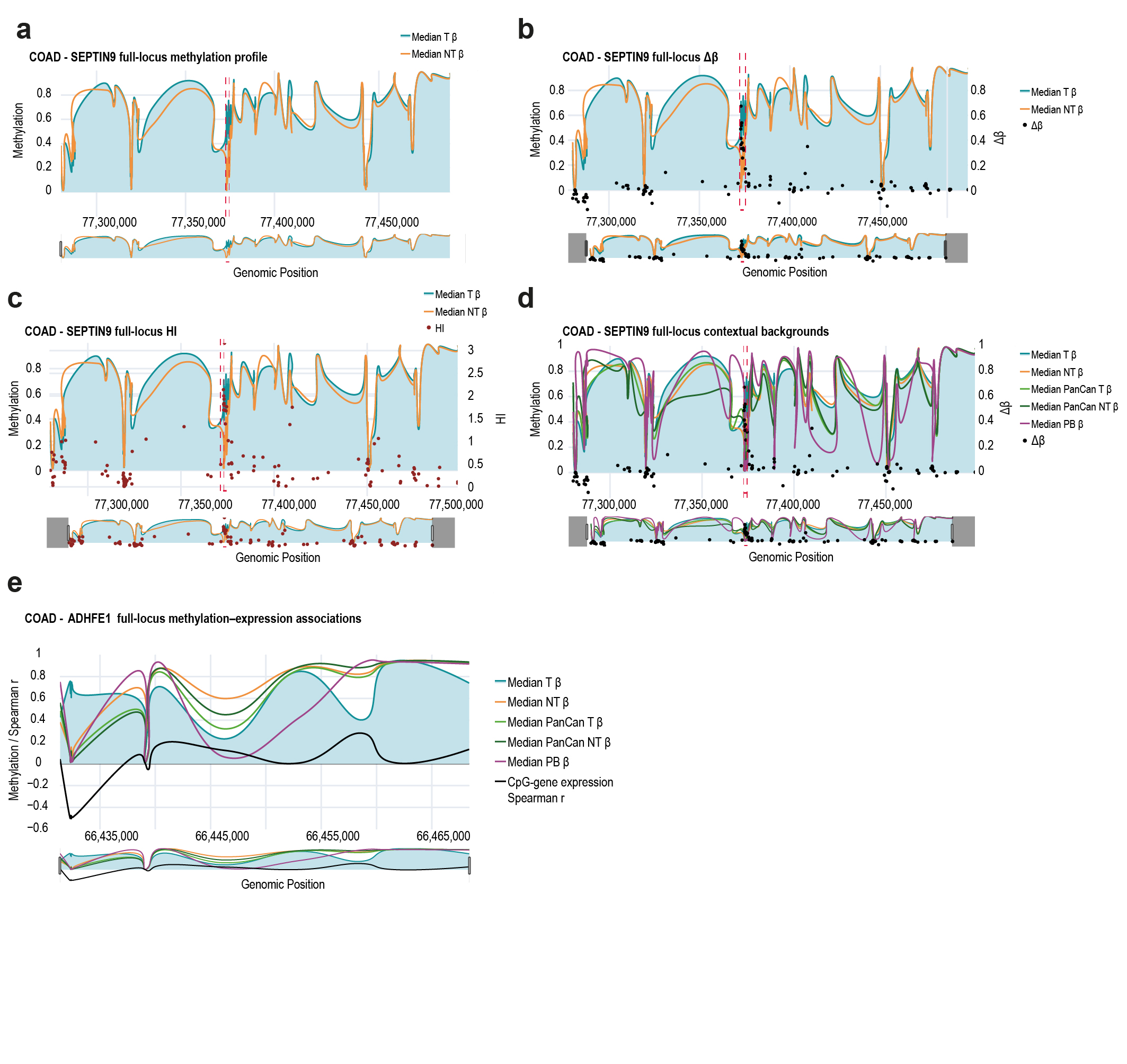

### Extended Data Fig 3

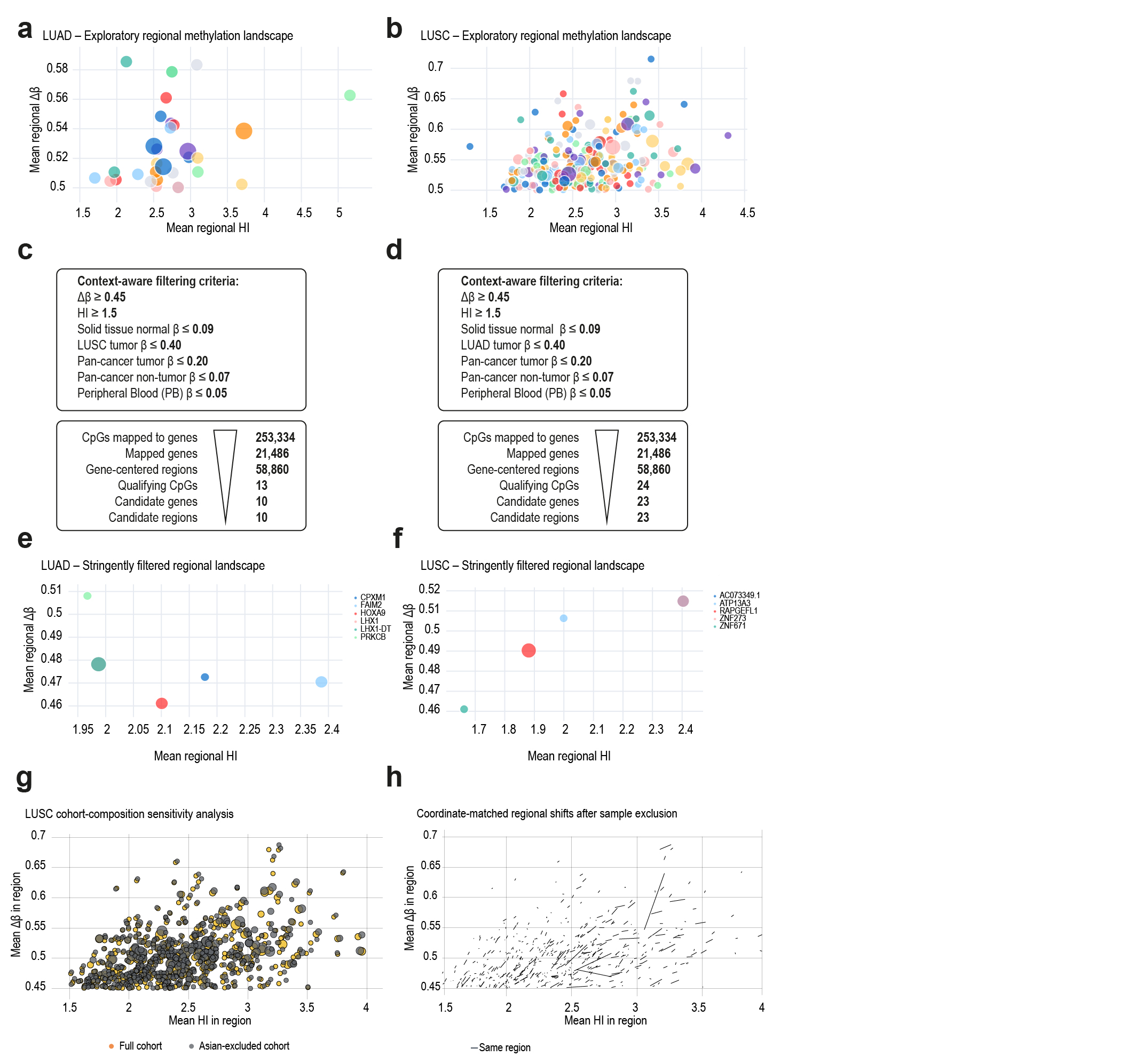
